# Comparing Approaches to Specimen Identification using Neotropical Freshwater Fishes in the Barra del Colorado Wildlife Refuge, Costa Rica

**DOI:** 10.1101/2023.10.02.560474

**Authors:** Taegan JM Perez, JP Fontenelle, Matthew A. Kolmann, Arturo Angulo, Hernán López-Fernández, Nathan R. Lovejoy

## Abstract

As global biodiversity declines continue, conservation efforts are increasingly important in megadiverse areas such as the Neotropics where biodiversity is especially imperiled. The accurate identification of specimens is critical to successful conservation plans. However, in groups such as freshwater fishes, different identification methodologies have documented challenges. Using a biodiversity survey of fishes from the Barra del Colorado Wildlife Refuge in northeastern Costa Rica, we compared: (1) morphological identifications in the field, (2) morphological identifications in the lab by experts, (3) DNA barcode-based identifications, and (4) identifications based on an integrative approach. Our results suggest that both barcode-based identifications and field morphological identifications provided fewer correct species identifications than lab identifications performed by experts using morphology. We attribute shortfalls of DNA barcoding in this case to the misidentification of reference material, the use of outdated taxonomy for references sequences, and the non-uniform representation of groups in public databases across taxa. We suggest the use of an integrative approach to identify freshwater fishes in Costa Rica and other megadiverse areas of the Neotropics where similar issues with public barcode reference libraries exist. We also recommend the creation of regional curated barcode reference libraries to aid in the identification of traditionally difficult to identify species/specimens. We also provide the most up to date species list for the ichthyofauna of the Barra del Colorado Wildlife Refuge identifying 51 species from 42 genera, 21 families, and 17 orders. Generating accurate species lists for protected areas and areas of importance will provide conservation practitioners with effective tools for tracking diversity changes over time.

## Introduction

The unprecedented rate of biodiversity loss due to anthropogenic actions brings attention to the urgent need for effective conservation initiatives. Prior to the establishment of initiatives, a better understanding of the overall health of the ecosystem in question is required. The health an ecosystem can be assessed by monitoring changes in species richness, diversity, and distribution over time (Shoo et al., 2006). The success of this measure is heavily dependent on the accurate identification of species (taxonomy), their biology and distribution patterns (Chovanec et al., 2003; Dudgeon et al., 2006). This complex scenario is aggravated by the taxonomic crisis (or impediment): a current lack of interest in taxonomy training and decreased financial incentives to study diversity (Pires & Marinoni, 2010; Ricciardi et al., 2020; Engel et al., 2021). This impediment is a key obstacle preventing the effective conservation of global biodiversity by confounding specimen identifications; restricting assessments of ecosystem health, and consequently limiting conservation initiatives (Sheth & Thaker, 2017; Ricciardi et al., 2020).

### Identification Methods

A critical first step in assessing ecosystem health is establishing a robust taxonomy of species present in a given area. Conventionally, taxonomic identification is morphology-based (Packer et al., 2009). Morphology-based identifications make use of different sets of quantitative and qualitative phenotypic characters, which include allometry, phylogenetics, morphometrics, and meristics (Hebert et al., 2003; Ward et al., 2009). For certain taxonomic groups, convergence in morphologically diagnostic characters and cryptic species pose a challenge for morphological-based identification (Pires & Marinoni, 2010; Marancik et al., 2020; Pereira et al., 2021). In addition, the ‘taxonomic impediment’: the small number of trained taxonomists and funding compared to the need for these specialists, complicates these issues further creating an intrinsic problem with incomplete/outdated references (Pires & Marinoni, 2010; Sheth & Thaker, 2017; Riccardi et al., 2020; Engel et al., 2021). These challenges can make identifying specimens/diversity difficult. Thus, in an effort to better resolve datasets identified by morphology, a complementary method: DNA barcoding has become increasingly popular for identification.

DNA barcoding is a technique for identifying specimens through the use of standardized genomic regions, called ‘barcoding genes’ (Ward et al., 2009). The mitochondrial gene Cytochrome Oxidase Subunit I (*coI*) is conventionally used for animals (Antil et al., 2022). Many studies have shown that *coI* sequences differ among closely related species to an extent that species identity can be established by comparing their sequences, even discriminating recently diverged cryptic species (Castresana et al., 1994; Hebert et al., 2003; Hubert et al., 2008; Ward et al., 2009; Lara et al., 2010; Pires & Marinoni, 2010; Kress et al., 2015; Creer et al., 2016).

One issue plaguing DNA barcoding as an identification technique is inconsistencies in identification of reference material. Ideally, once obtained, the barcode sequence of a specimen is compared to a public or private sequence database (or barcode reference library), produced from accurately identified specimens (Fort et al., 2021). Theoretically, these database sequences are produced from museum voucher specimens identified by experts (Sheth & Thaker, 2017; Buckner et al., 2021). However, ambiguous/erroneous sequence data added to public databases cause inconsistencies in identification and issues in downstream applications. (Grant et al., 2021; Rimet et al., 2021). Without properly identified voucher specimens to validate barcodes, the credibility of these barcodes and the associated public databases decreases (Rimet et al., 2021; Mulcahy et al., 2022). This negatively impacts the efficacy of DNA barcoding for estimating species diversity and subsequently, for conservation and management purposes.

While morphology- and DNA-based taxonomic approaches have demonstrated efficiency in species identification, both have caveats (Ward et al., 2005; Weigt et al., 2012; Ramirez et al., 2017; Pollack et al., 2018; Pentinsaari et al., 2020). Integrative taxonomic approaches that combine complementary datasets such as DNA and morphology, have proven to be efficient in addressing gaps left by each individual method (Breedy et al., 2012; Riedel et al., 2013; von Beeren et al., 2016; Duong et al., 2020; Zamani et al., 2022). The term ‘integrative taxonomy’ can be used to describe two processes, species delimitation/discovery and specimen identification (Pires & Marinoni, 2010; Pereira et al., 2021); herein we use the term integrative taxonomy to refer to specimen identification. The use of multiple data sources to support a specimen identity and subsequently the species presence in an area provides strong evidence that can be used for implementation of conservation plans, a key component to secure financial support for conservation (Sheth & Thaker, 2017). An integrative approach to specimen identification can also accelerate specimen identification by having morphology and barcoding work in tandem, in turn shortening the time frame for identifying key areas in need of protection (Pereira et al., 2021; Zamani et al., 2022).

### Study system: Costa Rica’s Barra del Colorado Wildlife Refuge

The Neotropics, arguably the most diverse ecoregion in the planet, is an important area for conservation efforts due to the rapid rate at which this diversity is disappearing (Antonelli, 2022). These high levels of diversity and endemism are related to the deep and complex evolutionary history of the species found there (Antonelli, 2022). Despite many efforts, Neotropical biodiversity is still underestimated (Antonelli et al., 2018). This is generally explained as due to a lack of taxonomic training, the uneven distribution of resources among taxa, and/or morphological complexity (Antonelli et al., 2018). This incomplete foundation is impractical for building effective conservation plans particularly when assessing ecosystem health using diversity proxy (Chovanec et al., 2003; Dudgeon et al., 2006). This is also complicated by the difficulty in producing accurate identifications for specimens. The Neotropics are susceptible to anthropogenic changes, and its freshwater systems are especially vulnerable (Albert et al., 2020). Currently, human activities are causing widespread habitat degradation (mostly due to agricultural expansion), and hydrological alterations (Albert et al., 2020). Additionally, the introduction of non-native species is also a major threat, as competition, predation, and disease all contribute to diversity declines (Albert et al., 2020).

Neotropical freshwater fishes represent the most species rich vertebrate fauna and new species discovered and described each year (Antonelli et al., 2018; Albert et al., 2020). Successfully conserving this diversity relies not only on protecting habitat, but also on understanding the taxonomy, biology, and distribution of Neotropical freshwater fish species. This megadiverse ichthyofauna typifies a challenging group for morphology-based identifications. Exceedingly high level of species diversity, deficient species descriptions and a high number of cryptic species and species complexes make hamper accurate species identification (Pereira et al., 2021). The use of DNA barcoding to circumvent limitations of morphology-based taxonomy in Neotropical fishes has proven to be an effective approach to further investigate diversity of this group. Previous studies have demonstrated the ability of *coI* to distinguish among species of freshwater fishes in different regions across the globe (Hubert et al., 2008; Lara et al., 2010; Pereira et al., 2013; Díaz et al., 2016; Shaw et al., 2016; Ali et al., 2020; Liu et al., 2020). However, DNA barcoding may be more challenging for Neotropical fishes given the reliance of this method on morphology for initial identification of voucher specimens and the known inconsistencies in public barcode databases. (Meier et al., 2008; Fort et al., 2021; Grant et al., 2021; Hammer et al., 2021). An integrative approach can aid in the identification of Neotropical freshwater fishes. The combination of data from morphology and barcoding can overcome the challenges each method presents independently (Gomes et al., 2015: Pugedo et al., 2016; Braga et al., 2021; Pereira et al., 2021).

The Barra del Colorado Wildlife Refuge was established in 1985 and extends from the San Juan River (Costa Rica’s border with Nicaragua) south to the northern tip of Tortuguero National Park (TNP) (Khazen et al., 2016). BCWR covers approximately 92,000 hectares of land, and in combination with TNP (19,000ha), makes up the Tortuguero Conservation Area (ACTo) (Grant et al., 2013). Established to link the Indo Maíz Reserve in southeastern Nicaragua with TNP, the combination of these areas forms the largest contiguous protected area of lowland Atlantic tropical wet rainforest in Central America, at approximately 111,000ha (Burger, 1997; Wickham, 2001; Khazen et al., 2016).

Several morphology-based species lists and diversity assessments for the fishes of both Costa Rica and Tortuguero region have been published (Angulo, 2021; Angulo et al., 2013; Bussing, 1998; Miller, 1966). Angulo et al. (2013) published an annotated checklist of the freshwater fishes in both continental and insular Costa Rica. This work records 250 species, adding 108 species to the most recent dichotomous key for the region: Bussing (1998). This checklist was recently updated by Angulo (2021) and provides the most recent checklist of the freshwater fishes in Costa Rica, listing 119 species in the “Tortuguero basin”, a region that includes ACTo. No comprehensive fish biodiversity study of BCWR has been completed.

### Goals

Given the issues associated with single data methods for identification (such as morphology or DNA barcoding), the potential of integrative approaches to aid in proper specimen identification, and that proper identification is essential in effectively understanding and documenting biodiversity, we investigated the effectiveness of different methods of specimen identification using a survey of fishes in the BCWR. Our study tests the effectiveness of DNA barcoding and morphological approaches, by comparing them to an integrative approach that we consider represents the most accurate method of specimen identification..

## Methods

### Fish Tissue Collection/Preservation

Fish specimens were collected biannually in February from 2013 to 2019 in the region surrounding Caño Palma Field Station in the Barra Del Colorado Wildlife Reserve, Costa Rica. A total of 10 sites were sampled in the four combined efforts (Figure 1). A list of sites per year, as well as locality coordinates, are presented in Table 1. Collection was done using a mix of seine nets, gill nets, dip nets, minnow nets, and electrofishing, with each collection effort involving six to 10 researchers collecting for approximately 1-2 hours. Fish were euthanized using clove oil solution following Keene et al. (1998). Samples of muscle tissue were excised from specimens and preserved in 95% ethanol solution, and subsequently stored in 95% ethanol solution at -20°C. For each site collected during each expedition, an attempt was made to sample tissues from two specimens per putative species. Euthanized fishes were preserved in a 10% formaldehyde solution in the field, and later transferred to 70% ethanol for long-term storage. Unfortunately, the 2017 collection of fishes was lost in the field; however, tissues from the 2017 expedition were used in this study.

**Figure 1.**
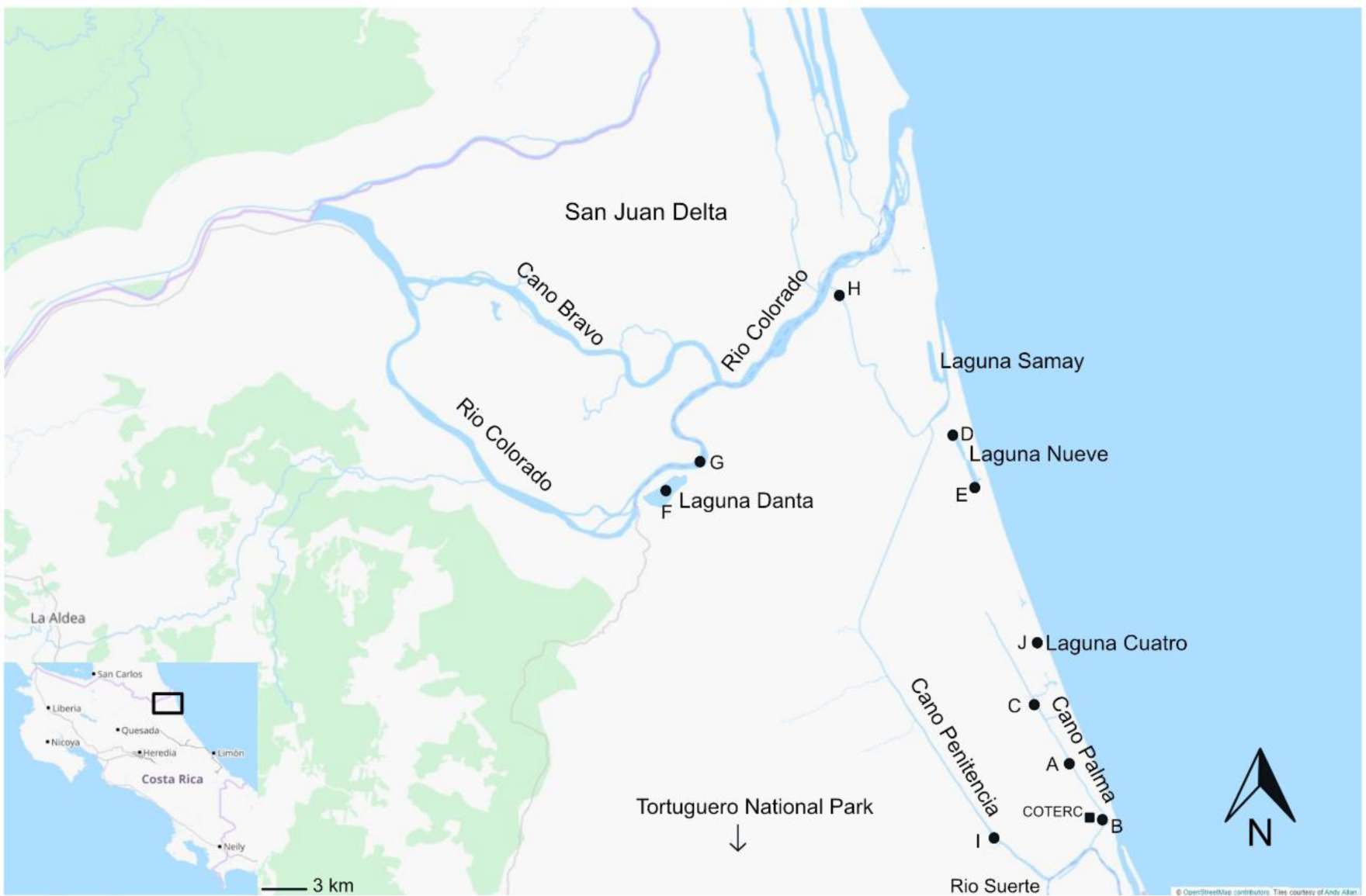
Map showing collection sites (black circles) visited over four collection expeditions from 2013-2019. The black square on the inset map shows the sites in relation to Costa Rica.

**Table 1.** List of sites visited over the four collection expeditions, including coordinates, site description, and years the sites were sampled.

| Site | Site Name | Latitude | Longitude | Description | 2013 | 2015 | 2017 | 2019 |
| --- | --- | --- | --- | --- | --- | --- | --- | --- |
| **A** | Small Streams entering Cano Palma | 10.6055 | -83.53412 | Small streams. Brown water, muddy bottom that varied from hard to soft mud along the site; debris in water included branches and leaves; murky and swampy with low visibility. Stream cover ranged from 65-100% cover. Stream width ranged from 2m-6m. Stream depth was 1-2.5m. Depth of capture for specimens ranged from 0m at minimum to 2.5m at maximum. Distance from shore that specimens were collected ranged from 0m at minimum to 600m at maximum along the site. Duration of collection ranged from 1hr to 2hr 4min along the site. Water temperature varied from 20.4-27.8C°. Water pH of 5.35. Air temperature varied from 25.5-29.5C°. O2 content varied from 566-1336ppm. Fishing methods included dip nets, seine nets, gill nets, minnow traps and electro fishing | x | x | x | x |
| **B** | Cano Palma | 10.5937 | -83.52725 | Black water canal. Muddy bottom, low visibility. Shores of river had *Rapphia* palms, *pentadetha*, and ferns. Woody material also floated on surface. Stream cover varied from 0-20%. Stream width was 20m. Depth of capture for specimens ranged from 0m at minimum to 0.5m at maximum. Distance from the shore that specimens were collected ranged from 1m at minimum to 10m at maximum. Water temperature varied from 23.6 to 28.5C°. Air temperature was 26C°. Conductivity was 1108us. Fishing method was active night fishing with dipnets from both the dock at COTERC and by boat. | x | x | x | x |
| **C** | Stream from Cano Palma to Turtle Beach Lodge | 10.6233 | -83.54462 | Small stream. Brown water, muddy, soft bottom, debris in water included fallen leaves and tree roots and branches; water very murky with low visibility. Stream cover was about 65%. Stream width 5m. Water depth ranged from 1-6m. Depth of capture of specimens ranged from 0m at the minimum to 1.2m at the maximum. Distance from the shore that specimens were collected ranged from 0m at the minimum to 3.5m at the maximum. Duration of collection was about 1 hour. Water temperature was about 27.5C°. Air temperature varied from 28-29.5C°. O2 contents ranged from 566ppm to 1336ppm. Fishing methods included dip nets and seine nets. |  |  | x | x |
| **D** | Stream connecting Laguna Nueve to XX | 10.7081 | -83.57519 | Small stream. Water appeared, green, white, and brown in different areas of site. Visibility varied from low to moderate. Bottom composition included sandy silty bottoms and varied hard mud and soft mud bottoms. Materials at site included leaves and logs. Grass, shrubs, and thick rainforest were seen around site. Varied wildlife was noted at the site including howler monkeys, kingfishers, and butterflies. Stream cover varied across the site, ranging from 30% to 70%. Stream width ranged from 4m to 10m. Stream depth ranged from 1.5m to 2m. Depth of capture for the specimens collected ranged from 0m at minimum to 1.8m. Distance from the shore that specimens were collected ranged from 0m at minimum to 100m at maximum. Duration of collection efforts varied from 52min to 2hr and 16min. Water temperature varied across the site from 23 to 31.4C° with pH between 6 and 6.4. Air temperature varied as well, ranging 26 to 30C°. Fishing methods included dip nets seine nets, gill nets, and electric fish finders. | x | x | x | x |
| **E** | Laguna Nueve | 10.6981 | -83.56882 | Lake. Greyish water with a silty, sandy bottom. Visibility was high. Grass and trees present on the shores. Depth of capture for specimens ranged from 0.1m at minimum to 1.5m at maximum. Distance from the shore that specimens were captured ranged from 0.05m at minimum to 115m at maximum. Water temperature was 30C° with a pH of 7. Fishing methods included dip nets and seine nets | x |  | x |  |
| **F** | Laguna Danta | 10.6939 | -83.65355 | Lake. Dark brown water with low visibility, sludgy bottom with detritus, thick grasses present on shore. No cover above the site, out in the open. No stream width noted. Water depth was approximately 1m. Depth of capture of specimens ranged 0.05m at minimum to 1m maximum. Distance from the shore ranged from 0.05m at minimum to 30m at maximum. The duration of the collection effort was 2hrs. Water temperature was 30C°. Air temperature was 35C°. Fishing methods at this site included dip nets and seine nets. | x |  |  |  |
| **G** | Rio Colorado | 10.6982 | -83.64732 | Large river. White waters with a sandy bottom. Visibility varied between years. Submerged grass and grassy islands in river. In 2013 the site contained a wide exposed beach due to a prolonged dry season. Downstream from Delta San Juan. Open water with no cover. Depth of small stream off of Rio Colorado was 0.5m. Depth of capture for specimens ranged from 0m at minimum to 1.5m at maximum. Distance from the shore that specimens were captured ranged from 0m at minimum to 1.5m at maximum. Duration of collection effort was 1hr. Water temperature was 34.8C° with a pH of 6.68. Air temperature was 27C°. Fishing methods included: seine nets. | x | x |  |  |
| **H** | Shore of river connecting Rio Colorado to Cano | 10.745 | -83.6053 | River. White waters with a silty mud bottom. Submerged grass present along bottom. Fish collected near the shore of Rio Colorado that connects Rio Colorado to Cano Penitencia. Stream cover not listed. Stream width was approximately 50m. The depth of capture for specimens ranged from 0m at minimum to 1.5m at maximum. Distance from the shore that specimens were collected ranged from 0m at minimum to 4m maximum. Duration of collection not noted. Water temperature was 25.5C° with a pH of 6.62. Air temperature was 30C°. Fishing methods included seine nets. |  | x |  |  |
| **I** | Suerte Cito | 10.5946 | -83.56641 | Stream. Green, brown waters with a muddy bottom that ranged from soft mud to moderately firm mud. Visibility was moderate. Mainly woody debris is water. Large trees and many green plants along the shore. Site is about 30minutes from the COTERC station. Stream cover varied between 40-50% cover. Stream width was approximately 10m. Stream depth was >2m. Depth of capture for specimens ranged from 0m at minimum to 1.6m at maximum. Distance from the shore that specimens were captured ranged from 0m at minimum to 5m t maximum. Duration of collection effort was 3hr 10min. Water temperature was 26.3C°. Air temperature was 25C°. Note that electric fish caught at this site were found in a small stream off of the main site, with very muddy water and bottom, and high turbidity. Fishing methods included: dip nets, seine nets, and electric fish finders |  |  | x | x |
| **J** | Laguna Cuatro | 10.6337 | -83.54392 | Lake. Brown water with a sandy bottom. Moderate to high visibility. Visible tree roots. Smaller grassy islands dispersed throughout lake. Nearby to Caribbean Sea beach. Stream cover was approximately 15%. Stream width was approximately 100m. Stream depth was not recorded. Depth of capture for specimens ranged from 0m at minimum to 1m at maximum. Distance from the shore that specimens were caught ranged from 0m at minimum to 50m at maximum. Duration of collection effort was 55min. Water temperature was 28.6C°. Air temperature was 29C°. Fishing methods included dip nets and seine nets. |  |  | x | x |

### Morphological Identification

In the field, before preservation, specimens were identified using morphology according to the keys provided in Bussing (1998). Identifications were primarily completed by undergraduate students, working with a graduate student teaching assistant and course instructor.

In some cases, specimens were identified based on morphology by taxonomic experts at natural history collections. Dr. Hernan Lopez-Fernandez, Royal Ontario Museum (now at University of Michigan), identified only the cichlid fishes collected during the 2013 and 2015 expeditions. Arturo Angulo, University of Costa Rica, identified all specimens collected during the 2019 expedition.

### DNA Barcode Identification

Genomic DNA was extracted from muscle tissue using the DNeasy Blood and Tissue Kit following the manufactures’ protocol (QIAGEN, Canada).

The mitochondrial barcoding gene, Cytochrome Oxidase subunit 1 (*coI*) was amplified using the primers COIFishF2 and COIFish R2 (Ward et al., 2005) (Table 2). Master mixes were created using the following: 13.75µl of ddH2O, 2.5µl of a combined buffer of 60% SO4 buffer and 40%KCl, 3.0µl of MgCl2 (25mM), 1.0µ of dNTP (10mM), 1.25µl of each primer, and 0.25µl of Taq Polymerase, totalling 23µl. A volume of 2µl of DNA was added to the master mix, accounting for a final volume of 25µl. Thermocycler conditions were as follows: 94°C for 2 minutes, 94°C for 30s, 52°C for 40s, 72°C for 1 minute and 30s. This series of steps was repeated 35 times in the cycler. Finally, PCR samples were set at 72°C for 5 minutes before resting at 4°C until removed from cycler. This protocol did not yield successful amplifications for each sample. I used a second combination of primers COIFishF1 and COIFishR1 primers (Ward et al., 2005), when amplification using the first combination of primers was unsuccessful (Table 2). When necessary, the volume of DNA per sample was increased to 3µl, removing 1µl ddH20 from the master mix. Each PCR run was accompanied by a negative control (22 or 23µl of master mix and 3 or 2µl of ddH2O respectively). PCR products were then checked for success by visualizing on Agarose Gel dyed with Red Safe®, where 5µl of PCR product was mixed with 2µl of 6x loading dye, a DNA ladder was added to visualize DNA lengths, and 2µl of 6x loading dye was added to 5µl Gels were then run for 30 minutes on 80 v and 80amps. PCR product was purified using a custom mix of Exonuclease I and Calf Intestinal Alkaline Phosphotase (EXO-CIAP). A volume of 4µl of EXO-CIAP was added to each PCR product vial to remove excess primers, nucleotides, and buffers. The PCR products were incubated in a thermocycler at 37°C for 30 minutes, 85°C for 15 minutes, and stored at 4°C. This protocol was adapted from Li et al. (1991), Hanke and Wink (1995), Nordström et al. (2000), Watanabe et al. (2010).

**Table 2.**
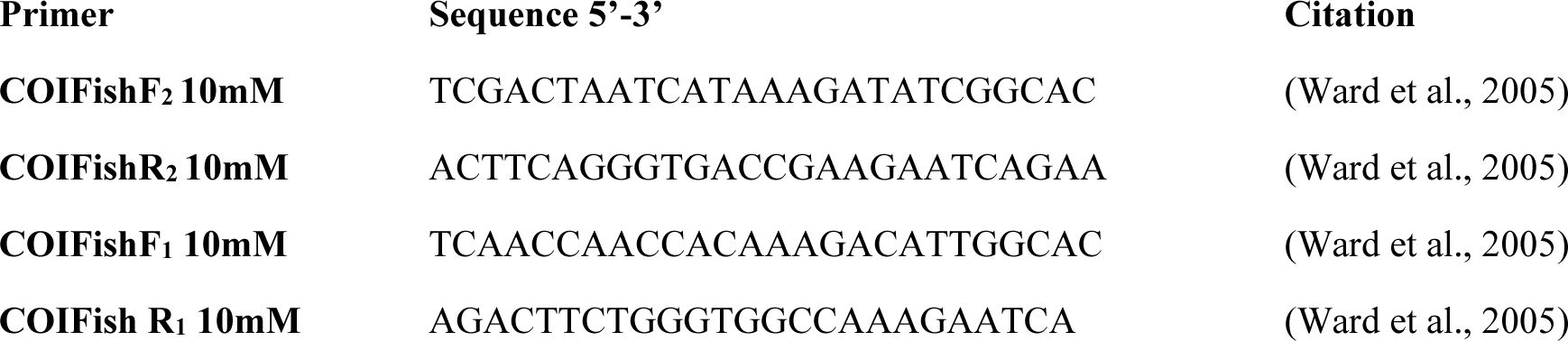
Primers used to amplify Cytochrome Oxidase Subunit 1 gene from Ward et al., (2005).

Purified PCR products were sequenced at the Centre for Applied Genomics: Peter Gilgan Centre for Research and Learning (Toronto, Ontario, Canada). Chromatographs were assembled, visually inspected, and manually edit in Geneious v9.8.1 (https://www.geneious.com). Consensus sequences were generated using Geneious’ default setting and aligned using the MUSCLE (Edgar, 2004a; Edgar, 2004b) plugin in Geneious.

Sequences of *coI* were exported from Geneious in FASTA file formats. Each sequence was submitted to Genbank’s BLAST search function and the Barcoding of Life Database’s (BOLD) search function using default search parameters (Benson, 2005; Ratnasingham & Hebert, 2007). For each sequence the three different species identifications with the highest percent similarity from each database from the top 100 results were recorded. The final barcode-based identification for each specimen was based on the highest similarity identification provided by the databases.

### Integrative Identification

For all specimens with *coI* sequences (239 total), identifications were also completed by integrating information from barcodes, field identifications, and expert identifications. This workflow is summarized in Figure 2. As the first step in this approach, working with each specimen individually, we queried the respective *coI* sequence in the databases Genbank and BOLD, producing a barcode identification. Then compared the morphological identification and the barcode identification. When expert or field identification and database identifications agreed, and when similarity of the specimen barcode was >98% to the database sequence(s), and if there were no ambiguities in the identification (i.e., the specimen barcode did not match multiple species in the databases), we accepted the identification. However, in cases where these criteria were not met, additional analyses were performed. For this project, an ambiguity was defined as a result that failed to provide a clear identification, for example, a specimen barcode that matched multiple species in the barcode databases.

**Figure 2.**
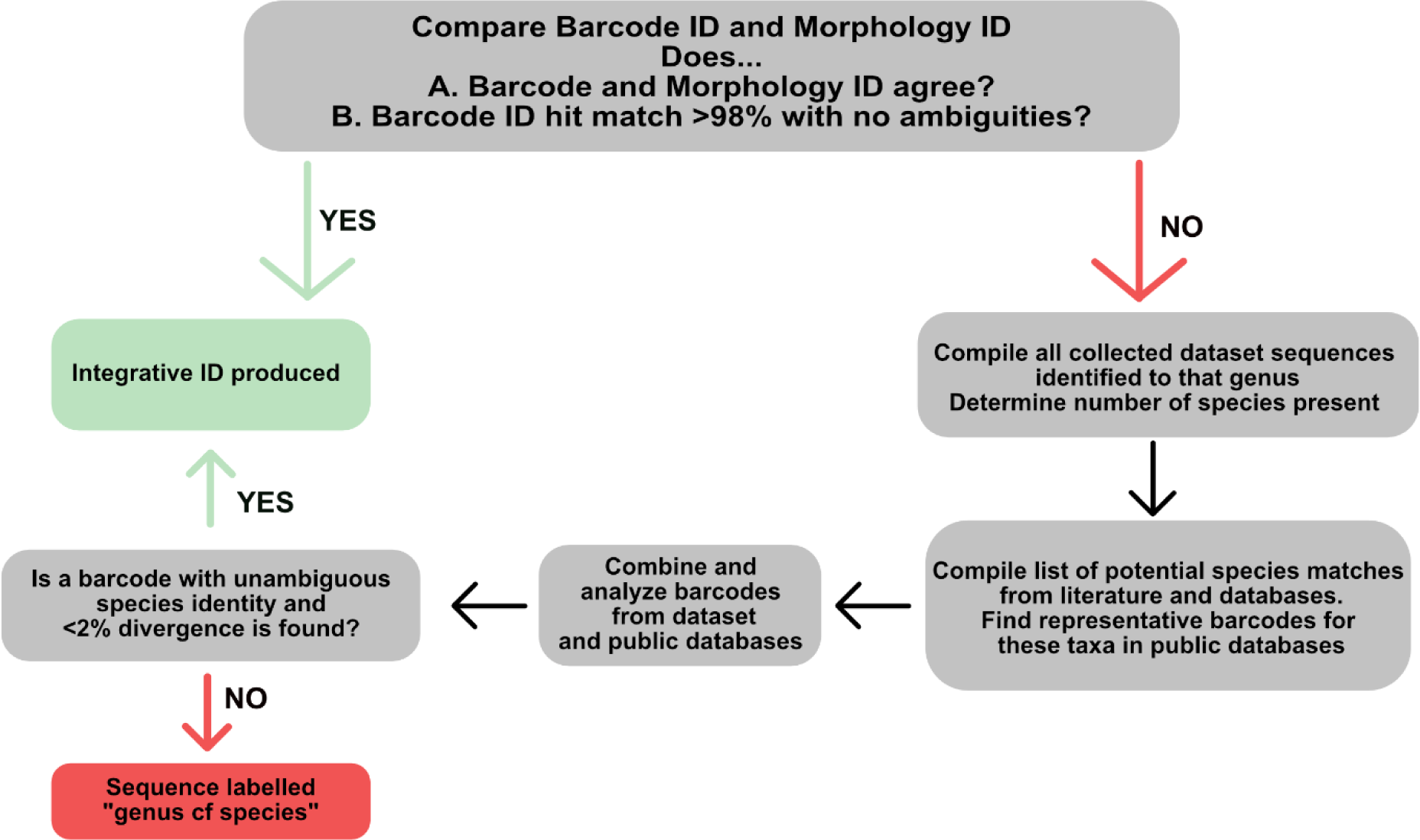
Workflow used to implement an integrative approach to the identification of specimens in BCWR.

For the additional analyses, we assembled and analyzed the barcodes of all BCWR specimens that belonged to a particular genus. And determined the genetic distances among all the specimens using the Kimura 2 parameter. This provided the information to determine the number of species represented, based on the idea that different species would be <2% diverged. Then, we compiled a list of species, genera, or families (in some cases) that might help to clarify the identity of our specimens. To determine which species, genera, or families should be considered, we reviewed the most recent diversity assessment for Costa Rican freshwater fishes by Angulo (2021), focusing on those closely related to the database results. We also reviewed the results from the queries in Genbank and BOLD and included the species that these databases identified. Once compiled, we searched Genbank and BOLD and downloaded barcodes for any of the species on the list, then determined the divergence between our specimens and all retrieved barcodes. The barcode match with the lowest divergence was used as the identification for the specimen. In most cases this resolved inconsistencies between the morphological and barcode identification. In some cases, a barcode from an unidentified specimen matched a barcode for a specimen with an expert morphological identification, and this allowed species identity to be established. In cases where the expert morphological identification and the barcode identification still did not agree, we utilized the expert identification. These identifications are listed as genus cf. species where the cf represents the uncertainty of the identification without additional data. The placements of barcode sequences in the neighbor-joining tree support the species identity assigned to them as the barcode sequences group with conspecifics.

### Holistic Approach to Determining Species Presence/Absence in BCWR

For the identification of individual specimens, we followed the approaches described above. However, this project also attempted to determine which species are present in BCWR using all available evidence, which we term a “holistic approach”. For this, we included species indicated by the expert identification of physical specimens, sequence identification (in cases where there are no physical specimens and where there were no inconsistencies in the database’s identification), unambiguous identification from photographs, unambiguous observations from locals, and species identified from the integrative approach previously described. Here, we use “unambiguous” to refer to evidence of species that would be highly unlikely to be confused with other species.

## Results

### Morphology-based Identification

Morphological identifications performed in the field using the keys in Bussing (1998) assigned the specimens collected in BCWR to 61 species.

Morphological identifications performed by curators of ichthyology at the University of Costa Rica and Royal Ontario Museum identified a total of 29 species. These identifications were based on cichlid fishes collected in 2013/2015, and all the specimens collected in 2019.

### Barcode-based identification

We analyzed a total of 239 sequences from specimens collected in BCWR. The results of the top 3 species matches for each specimen in the BCWR collection for each database can be found in Supplementary Material. The barcode approach assigned specimens to 51 different species using the Genbank database and 49 different species using the BOLD database.

Of the 239 specimens and associated sequences that were identified using the search functions in the databases, 93 were identified with high percent similarity matches (>98%) to one species. This included 19 species (*Dajaus monticola*, *Amatitlania siquia* (*nigrofasciata*), *Alfaro cultratus*, *Archocentrus centrarchus*, *Belonesox belizanus*, *Brycon costaricensis* (*guatemalensis*), *Cribroheros longimanus*, *Cribroheros rostratus*, *Ctenogobius fasciatus*, *Cynodonichthys isthmensis*, *Dormitator maculatus*, *Herotilapia multispinosa*, *Hyphessobrycon tortuguerae*, *Parachromis dovii*, *Parachromis friedrichsthalii* (*loisellei*), *Parachromis managuensis*, *Phallichthyes amates*, *Roeboides bouchellei* and *Vieja maculicauda*). The remaining 147 specimens could not be identified to species level due to match similarity below 98% or high similarity matches (>98%) to multiple species. Both cases prevented a clear identification for the queried sequences.

### Integrative identification

Our integrative approach assigned 239 specimens to 42 species (Table 3). Six specimens could only be identified to the genus level using this method: Hypostomus sp Invasive (N10526, N10528) and Anchoviella sp 1 (N13542, N13543, N13631, N13635). Using the integrative approach included 6 species that were only determined using this method (Centropomus cf. ensiferus, Centropomus pectinatus, Eleotris cf perniger, Mugil cf. hospes, Parachromis dovii, Achirus cf. lineatus).

**Table 3.**
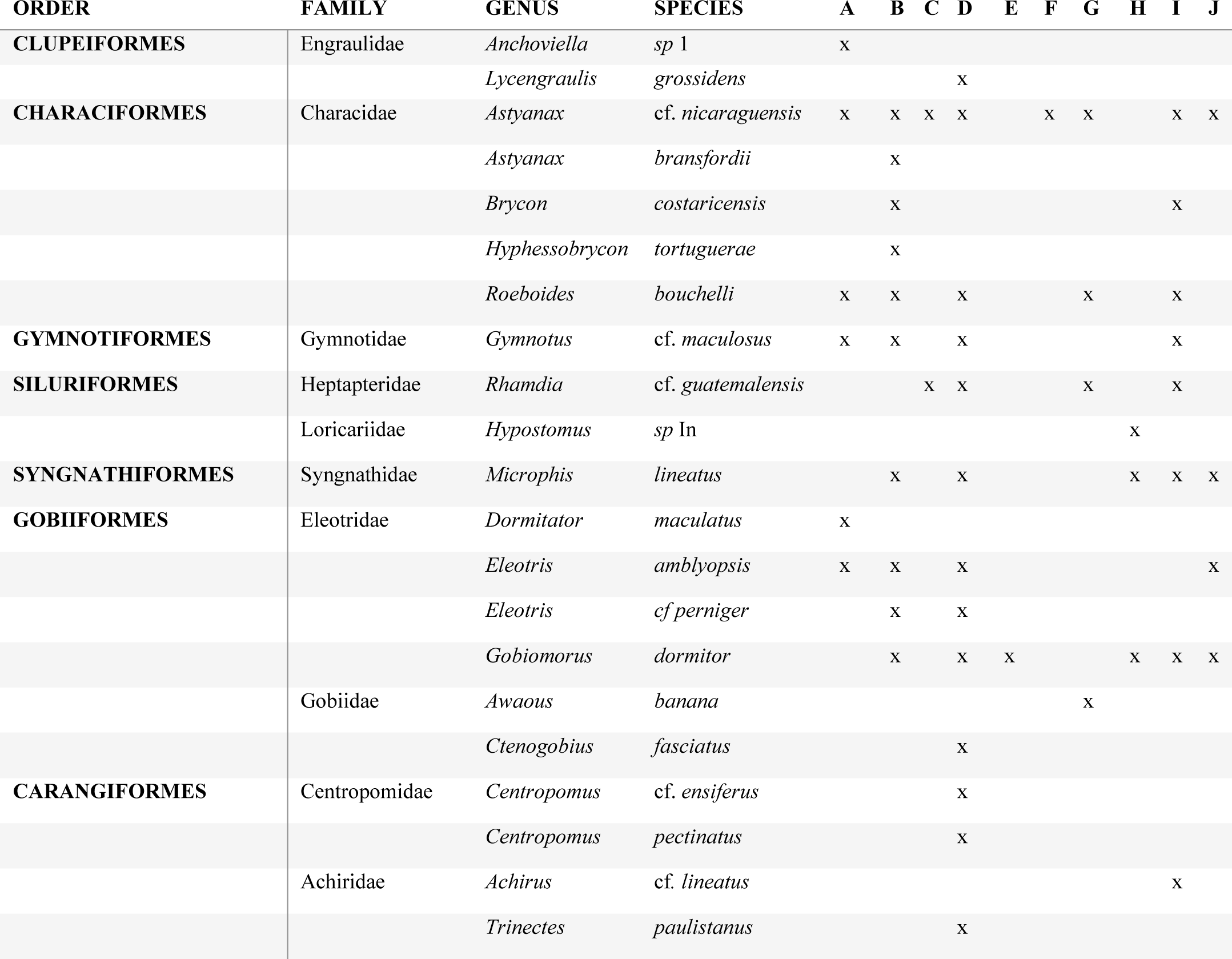
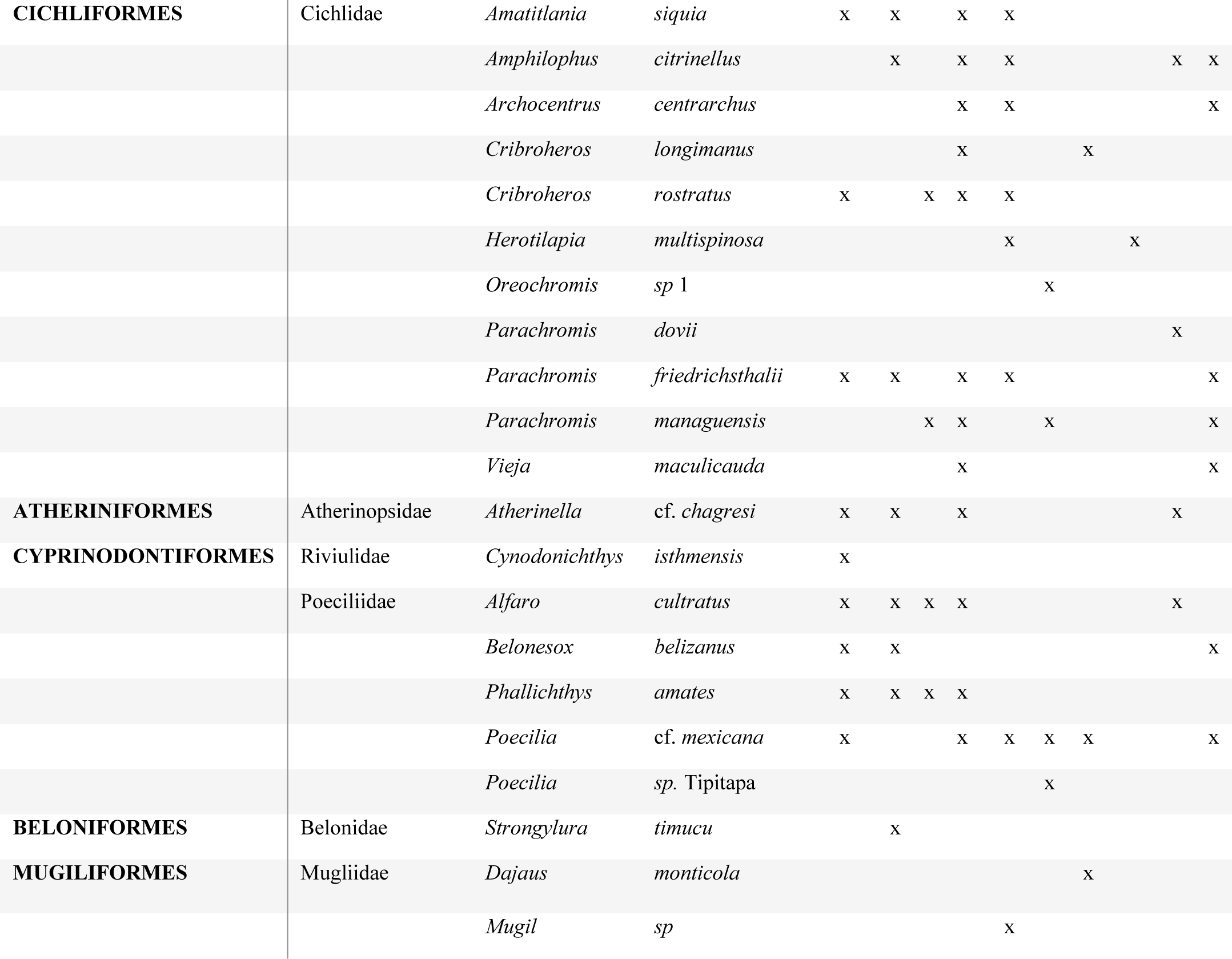
Freshwater fish species identified in BCWR using an integrative approach. Location of specimen collection also listed.

### Holistic species list for BCWR

The holistic approach to documenting species presence, based on evidence from multiple sources of data, found 51 species from 42 genera, 21 families, and 17 orders in BCWR (Table 4).

**Table 4.**
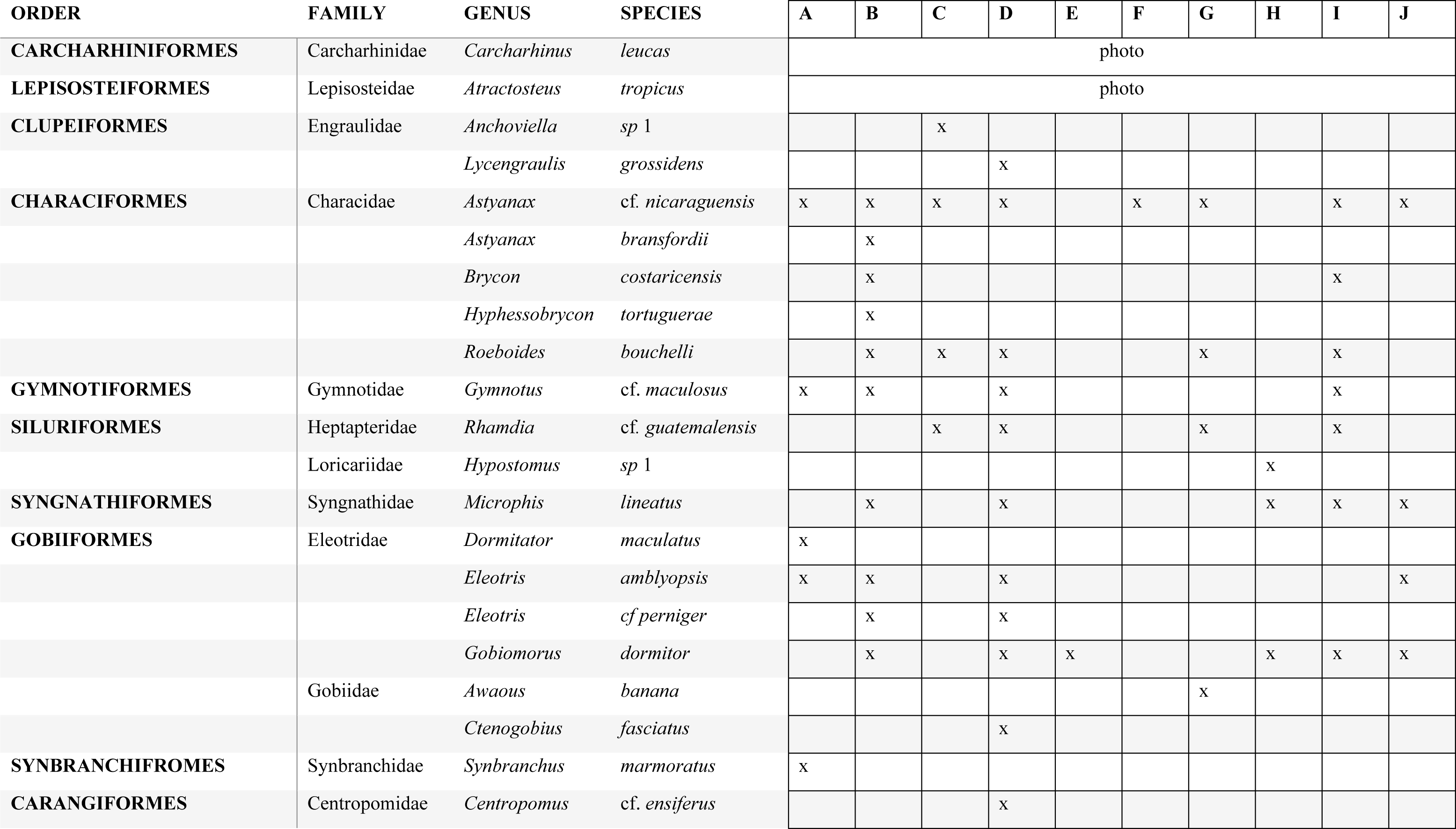
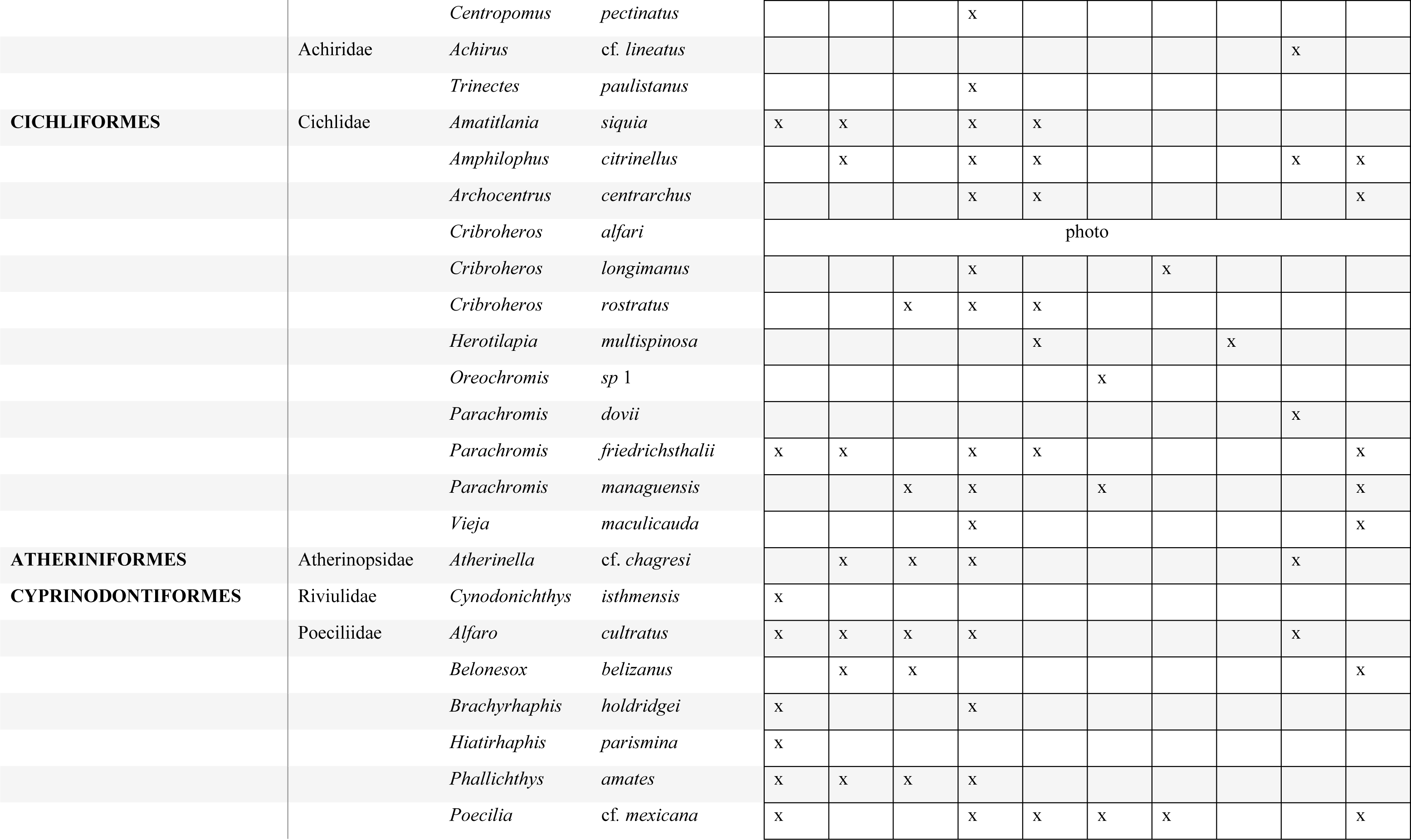
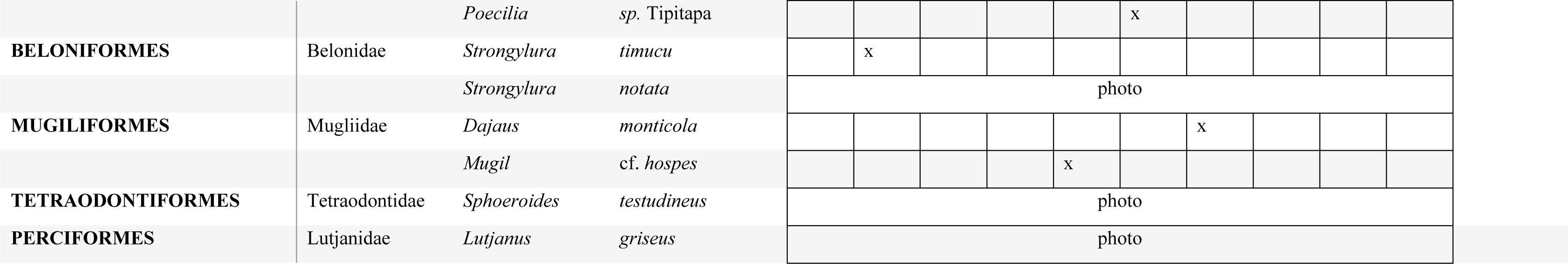
Species identified in BCWR using a holistic approach.

### Comparison of Identification Methods

To compare the different approaches to specimen identification, the 68 specimens that had field-based identifications, expert-based identifications, and barcode-based identifications from both databases were compared to identifications provided by the integrative approach (Figure 3). Since the specimens had been lost, none from the 2017 expedition were included in this analysis. We assumed that the integrative approach provides the best identification for a given specimen. Expert-based identifications produced the same identifications as the integrative approach more often than the barcode-based or field-based identifications. Interestingly, the barcode-based identifications and the field-based identifications both produced species level identifications that matched the integrative approach nearly 50% of the time (field-based 48.53%, BOLD database 54.41% and Genbank 51.47%). Identifications by experts using morphology provided the same genus level identification as the integrative approach nearly 100% of the time, whereas the barcode-based approach matched nearly 90% of the time (86.76 for BOLD, 91.18 for Genbank), and field-based was nearly 71% (70.59%).

**Figure 3.**
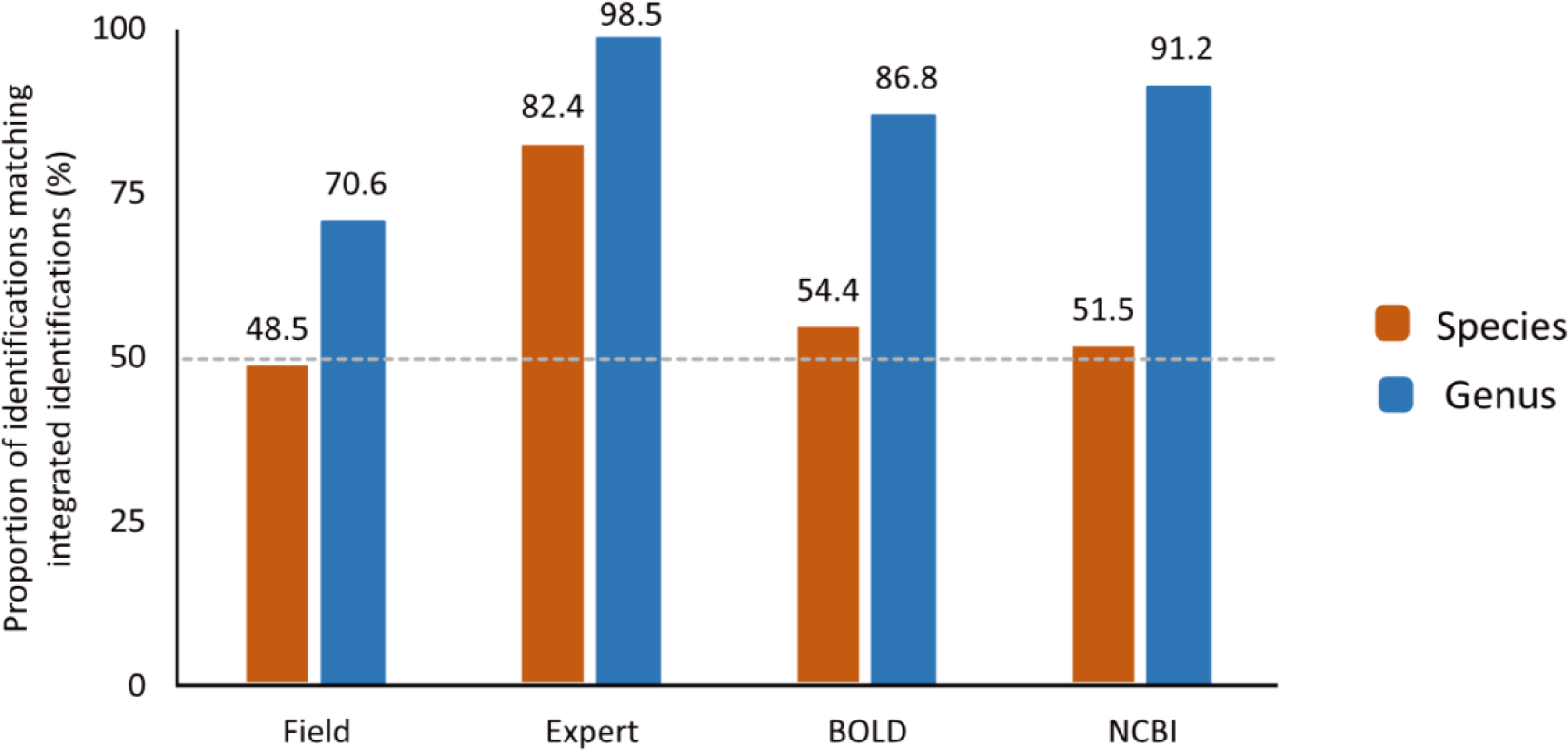
Comparison of 68 specimens identified using three methods to an integrative approach.

We also assessed the effectiveness of identification methods by comparing the identifications for a larger pool of 239 specimens that had field-based IDs, barcode-based IDs, and integrative approach-based IDs (Figure 4). As above, we compared the field and barcode IDs to integrative-based IDs. As observed in the previous comparison, overall species identifications based on both field and barcode approaches are less accurate compared to the integrative approach. The field-based approach performed similarly in this comparison as it did in the pervious. Field-based IDs matched the integrative approach less than 50% (40.83%) of the time at the species level and less than 75% (72.92%) of the time at the genus level. As seen in the previous comparison, the field-based method and the barcode-based method had a similar success rate at the species level, where all three methods produced the same identification as the integrative approach less than 50% of the time (field-based 40.83%, BOLD 42.08%, and Genbank 43.75%). The barcode-based approaches were more successful at identifying specimens to the genus level than the field-based identifications.

**Figure 4.**
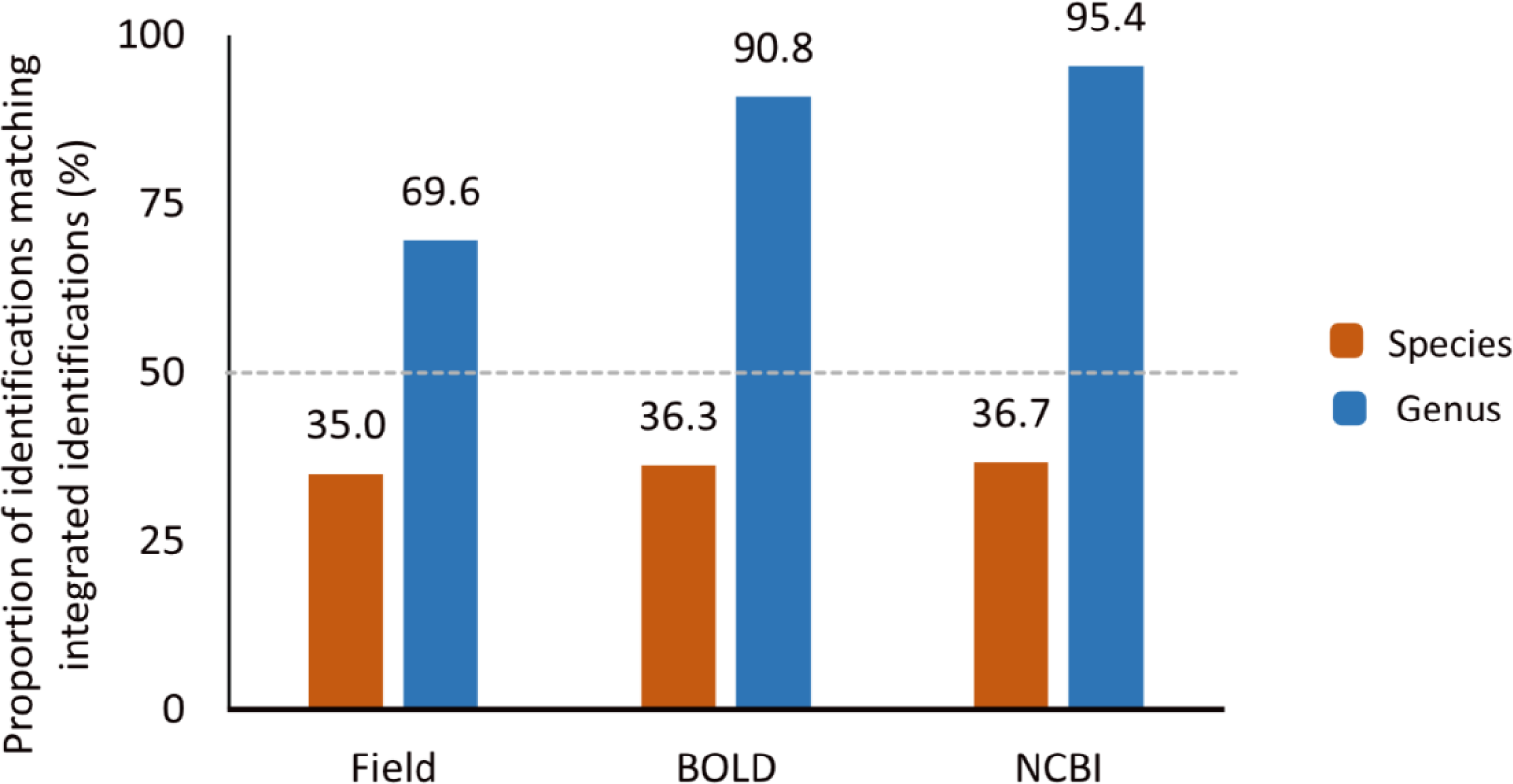
Comparison of 239 specimens identified using two methods to an integrative approach.

## Discussion

Integrative approaches are increasing in popularity to answer a wide range of biological questions (Will et al., 2005; Gomes et al., 2015; Hernández-Mena et al., 2019; Duong et al., 2020). One use for an integrative approach is in the identification of specimens (Pires & Marinoni, 2010; Sheth & Thaker, 2017). By integrating different data sources, we can provide more robust specimen identifications (Pires & Marinoni, 2010). We wanted to compare the use of morphology and DNA barcoding to an integrative approach to specimen identification in this region. DNA barcoding was of particular interest since no previous studies had assessed the diversity of freshwater fishes in Costa Rica using a molecular approach.

### Identification Methods

#### a. Morphology

Morphology is the primary resource for identification of specimens and taxonomy. In hyper diverse areas like the neotropics, morphological identification can be especially difficult for non-experts and is even more so in hyper diverse groups such as freshwater fishes whose taxonomy is also sometimes not fully understood (Schmitter-Soto, 2017). For this project, we examined and assessed two types of morphological identifications: field-based and expert-based. The field-based identifications were accomplished by students using a dichotomous key for the region by Bussing (1998). This key represents the most recent and accessible tool for non experts interested in identifying fishes in Costa Rica.

The identifications given following the Bussing (1998) key were the least accurate when compared to the integrative approach (Figure 3 and Figure 4). When compared to the integrative approach, the identification done in the field using morphology produced the same species level identification between 40% and 48% of the time (Figure 3 and Figure 4). This result is unsurprising given that more recent work in Costa Rica has added 108 new species that are not included in Bussing (1998) (Angulo et al., 2013). In addition, the field-based identifications were completed by students as part of an undergraduate course and were likely subject to more errors given their limited experience. This indicates that improvements to field-based identifications could derive from improvements to the Bussing (1998) key, or from additional training for the identifiers.

We also wanted to determine the success rate of expert-based morphological identifications. In this case, experts chosen were curators of fishes from natural history museums. Compared to field-based and barcode-based identifications, expert-based identifications were the most successful (Figure 3). Expert morphology-based identifications matched the integrative identifications for specimens at the species level 82% of the time (Figure 3). Dr. Angulo’s work on the 2019 specimens provides the most direct comparisons since all the fish specimens collected that year were identified. These results show that expert-based identifications are more successful than barcode-based and field identifications at both the species and genus level for this region.

However, experts are not always available for identification. As mentioned earlier, the taxonomic impediment is one of the biggest issues facing morphology-based identifications (Sheth & Thaker, 2017; Ricciardi et al., 2020). The number of scientists entering taxonomy and funding is decreasing, which is problematic as these specialists are needed for successful conservation of biodiversity (Ricciardi et al., 2020). Our results highlight how an integrative approach can efficiently guide specialists. The specimens that were not given the same identification for the integrative approach and expert-based identifications included specimens of *Strongylura* and *Centropomus.* In the case of *Strongylura*, a morphological reassessment, prompted by the DNA barcode analysis, indicated that the species identity was different than initially determined based on morphology.

Further analysis showed that morphology and DNA barcodes agreed that the specimens were *Strongylura timucu*. Upon first glance, the specimens of *Centropomus* appeared to represent a single species, however molecular data suggested the presence of two species in BCWR. When brought to experts’ attention, the specimens were reassessed, identifications revised, and the presence of two species of *Centropomus* was confirmed.

This study demonstrates the ability of an integrative approach to work in tandem with taxonomy, and the ability of this method to highlight specimens for special attention and areas for further research. Issues arising from the taxonomic impediment are typically cited as resulting from lack of funding and interest, however when specialists do exist, they are considered the best resource for taxonomy and identification (Pires & Marinoni, 2010). While DNA barcoding may be more accessible for non-experts, currently available public barcode libraries are not adequate, and specialists are needed to identify specimens in BCWR to produce foundations for biomonitoring and conservation.

#### b. Barcoding

For the specimens collected in BCWR, DNA barcoding did not identify all specimens. When looking at the 68 specimens that had been identified using all identification methods (field-based, expert identified in the lab, barcoded using Genbank, and BOLD), the barcode-based identifications from both databases produced the same identification as the integrative approach less than 50% of the time at the species level (Figure 3). Interestingly, the success of the barcode-based identifications was comparable to the field-based identifications performed by non-experts using a key that was identified as deficient. The databases were however able to successfully identify specimens to the genus level more than 85% of the time (Figure 3). While this provides some information about the diversity of freshwater fishes in BCWR, it does not provide the adequate detailed information authors have described as necessary for conservation of biodiversity (Dudgeon et al., 2006). Not all specimens used in this project had expert-based identifications; however, we could still compare the identifications for the 239 specimens with identifications that were field-based, barcoded using Genbank, and barcoded using BOLD (Figure 4). This comparison shows similar results, where the success rate of barcode identification using both databases was less than 50% and was comparable to the success rate of field-based identifications (Figure 4).

Why is the barcoding identification approach so unsuccessful for BCWR fishes? When using the barcode databases, we observed multiple instances where the databases were inconsistent in their designation of a species match (providing multiple species as matches) or were unable to provide a close match. These difficulties had several causes, as discussed below.

In some cases, we found that specimens/barcodes in the reference libraries were likely misidentified, leading to inconsistencies in query species identifications. The genus *Gobiomorus* provides a good example. Barcode-based identifications of specimens from BCWR indicated the species *Gobiomorus dormitor* and *Gobiomorus maculatus* both as matches. When conducting additional analyses on this group, we found that there were two phylogenetically distinct groups of barcodes within the public data sequences, likely representing *Gobiomorus dormitor* and *Gobiomorus maculatus*. However, within each of these two groups there were sequences listed as both *Gobiomorus dormitor* and *Gobiomorus maculatus*. A better understanding of the ecology and distributions of the two species was needed to determine a likely identification for these specimens. Authors have described species pairs existing on either side of a geographic barrier within Neotropical gobioids (Angulo, 2021; Thacker and Hardman, 2005) Within each pair, one species occurred in westward flowing Pacific drainages and the other in eastward flowing Atlantic drainages (Thacker & Hardman, 2005). Costa Rica shares a similar geologic pattern and there are distinct assemblages of fishes on either side of the Cordillera (Angulo, 2021; Angulo et al., 2013). Angulo (2021) confirms this pattern within the species pair *Gobiomorus maculatus* and *Gobiomorus dormitor*. This information strongly suggests that the two phylogenetically distinct groups of sequences in the barcode databases correspond to the two different species *Gobiomorus dormitor* and *Gobiomorus maculatus* but that samples within each group have been misidentified. Thus, although differentiating between these two species using barcode databases should be relatively straightforward, sample misidentifications mean this is not the case. Database barcode misidentifications can create a “snowball” effect, as samples erroneously identified are themselves entered in the databases leading to more errors in later projects.

We found that some species in the barcode databases, while not necessarily misidentified, were labelled with outdated taxonomic names. An example is the Cichlidae species *Vieja maculicauda*. There are sequences in both databases listed as *Paraneetroplus maculicauda*, a genus assignment now considered invalid assignment for this species (McMahan et al., 2015). This situation is not intuitive, particularly to those who may not have a deep understanding of the taxonomy of this group. The decreased accessibility of these databases because of a requirement for deeper taxonomic understanding to sift through data such as this is problematic if the goal is to be resource for non-experts and for conservation (Bianchi & Gonçalves, 2021). Bianchi and Gonçalves (2021) found that within the BOLD database on average 10% of misidentifications for a given taxon could be attributed to misspelling and invalid taxonomy, compromising the integrity of the database.

In many cases, there was incomplete representation of species in the barcode reference libraries. A particularly clear example of this was seen in the family Engraulidae (anchovies). Initial attempts to identify these specimens using the databases default search functions did not yield any matches above 98% similarity. Species that would be likely matches based on distribution and experts were not represented in the databases. This was also the case at the genus level. Identification of the anchovy specimens was reliant on only morphological identification. This was also not a singular occurrence in this study; inadequate representation in public databases prevented clear identification in other genera as well. Inadequate species representation prevents both direct identification of a specimen and creates data deficiencies in additional analyses. In addition, without this representation it also becomes impossible to corroborate a morphological identification with other data sources such as molecular data.

The records for individual sequences in BOLD are more complete than Genbank most of the time. However, BOLD contains significant amounts of privatised data. These records are not available to the public, meaning one cannot check the vouchers’ origin, the authors, locations, and additional details. This type of data is included in the repositories that BOLD uses to determine the identity of a queried sequence, making it more difficult for users to have confidence in identifications. Genbank generally had fewer complete records than BOLD but does not have privatised data meaning a user can access the records and determine which is most reliable when using the search functions to identify their specimens. When comparing the different identification methods to the integrative approach, we also compared the identifications given by each database to each other. We assumed that the two databases would provide the same identification for the same specimen since data is shared between them, however this only occurred 80% of the time for species level identifications. This suggests that there are content differences between the two databases that can result in each database producing a different species identification.

Overall, we found that DNA barcoding is insufficient for identifying most specimens collected in BCWR. The low success rate of DNA barcoding for this assemblage can be attributed to the problems discussed above with current public barcode libraries. Other authors have noted similar issues with public barcode databases, highlighting the need for reviews of this data (Diaz et al., 2016; Oliveira et al., 2016; Cariani et al., 2017; Bianchi & Gonçalves, 2021; Fort et al., 2021). Authors have suggested one way to circumvent these issues is to be selective in the public data used for specimen identification (Diaz et al., 2016; Cariani et al., 2017; Oliveira et al., 2016). Building barcode reference libraries for a group or region of interest has become increasingly popular where authors create protocols for exhaustively analyzing public data to include the best public data available (ex ELASMOMED DNA Barcode Reference Library from Cariani et al (2017)) (Diaz et al., 2016; Cariani et al., 2017; Oliveira et al., 2016; Fort et al., 2021; Antil et al., 2022).

### Implications, Next Steps, Recommendations

As mentioned, accurate identification of specimens is fundamental to diversity assessments and conservation more broadly. Currently, there is no resource for a non-expert that reliably identifies specimens found in BCWR. In addition, the inability of single data approaches to agree on an identification for a given specimen adds to confusion surrounding the diversity of freshwater fishes here. Without a solid understanding of the diversity of fishes in BCWR, effective conservation planning cannot occur. Moreover, the ability to monitor presence and absence of non-native species is severely impacted by the inability to identify specimens.

There are adjustments that can be made to current public DNA barcode libraries that would improve their effectiveness. The implementation of a reminder system for contributors, asking for them to check the sequences they have uploaded and confirm if the identification remained the best identification would help users, especially if the date the record was updated is displayed. In addition, the deeper involvement of experts in the vetting of records would help to correct errors of misidentifications or outdated taxonomic assignments.

The most difficult aspect of public DNA barcodes libraries as they currently stand is the feasibility of their use by non-experts. At the current time, especially for understudied regions like Costa Rica, public DNA barcode libraries are no easier to use than morphology due to the errors and underrepresentation of these libraries. One popular solution to the limitations of large public DNA barcode databases is the implementation of curated barcode reference libraries (Astrin et al., 2022). While suggested frameworks for the creation and implementation of the tools exist, significant funds are required at the inception of these projects (Astrin et al., 2022). For underfunded, understudied regions such as Costa Rica, the upfront cost associated with this solution becomes a formidable challenge. For Costa Rica, we suggest developing a framework that allows the best public data available to be found and exported the create the basis of a curated barcode reference library that can be built upon over time, as funds and information become available.

Attempting to fix these errors and improve the efficiency of these databases will take both time and effort however, the potential application of a great database is well known (Astrin et al., 2022).

### Conclusions

The protection of biodiversity continues to be at the forefront of science. Case studies such as these highlight areas that despite the known incredible diversity present, are still relatively understudied. In addition, for groups such as Neotropical freshwater fish there is still work that needs to be done concerning methods of specimen identification such that diversity can be better understood.

